# Curli Carrier Burden: a quantitative trait-level microbiome index for amyloidogenic bacterial signals in Parkinson’s disease gut metagenomes

**DOI:** 10.64898/2026.05.25.727557

**Authors:** Nabanita Ghosh, Krishnendu Sinha

## Abstract

**Background:** Parkinson’s disease (PD) gut metagenomic studies have repeatedly reported disease-associated shifts in microbial taxa, genes, and pathways. However, the field still lacks transparent trait-level indices that summarize biologically coherent microbial exposures. Curli fibres are extracellular bacterial amyloids produced by several Enterobacteriaceae and related taxa, and they provide a plausible microbiological bridge between gut microbial ecology, epithelial/immune interfaces, and alpha-synuclein-centered gut-brain-axis hypotheses. We introduced *Curli Carrier Burden* (CCB), a mathematically explicit, taxon-informed index that estimates the aggregate abundance of curated curli-carrier bacterial taxa in processed metagenomic profiles.

**Methods:** A curated curli-carrier candidate panel was converted into an evidence-weighted taxon set. For sample *s*, CCB was defined as 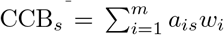, where *a*_*is*_ is the processed relative abundance of matched curli-carrier taxon *i* and *w*_*i*_ is an evidence weight reflecting curli-carrier confidence. We evaluated CCB in five main PD gut metagenomic evidence streams: Wallen 2022, Integrated-US, Mao Central China, Romano non-Wallen, and DuruIC 2024. Results were interpreted cohort-wise rather than as a formal meta-analysis.

**Results:** The CCB framework generated a reproducible sample-level microbial trait variable and enabled cohort-wise comparison of amyloidogenic bacterial burden. Wallen showed discovery-stage PD-associated elevation (724 samples; 31 matched curli taxa; Mann-Whitney *p* = 0.0020). Integrated-US provided supportive independent evidence (244 samples; 18 matched taxa; *p* = 0.0079). Mao Central China and DuruIC 2024 showed the same PD-greater-than-control direction by mean and median CCB, although their individual comparisons were not nominally significant. Romano non-Wallen provided a large multi-study analysis (600 samples; 29 matched curli-associated mOTUs taxa), with higher PD mean and median CCB in pooled analysis (*p* = 0.0036, Cliff’s *δ* = 0.137) and cohort-sensitive behavior under study-stratified permutation (*p* = 0.1974). Additional processed-cohort checks indicated that CCB interpretability depends on taxonomic representation and matched curli-candidate coverage, reinforcing the value of explicit compatibility reporting.

**Conclusions:** CCB is a novel, extensible, microbiology-informed index for quantifying amyloidogenic curli-carrier bacterial burden in processed gut metagenomic profiles. The current results support CCB as a useful exploratory trait-level variable for PD microbiome research and provide a principled route toward future raw-read, csg-operon, strain-resolved, and phenotype-aware studies of the curli-vagal PD axis.

## Background

Parkinson’s disease is increasingly studied as a systemic disorder in which gastrointestinal biology, microbial ecology, immune signaling, and the gut-brain axis may interact with neurodegenerative processes. Gut metagenomic studies have reported recurrent but cohort-dependent taxonomic and functional differences between PD and control groups (Romano et al., 2021; Wallen et al., 2022; Nishiwaki et al., 2024). These studies have made it clear that single-taxon associations are often sensitive to geography, study design, medication exposure, sequencing platform, and bioinformatic representation. Therefore, a complementary strategy is needed: rather than asking only whether one species is enriched or depleted, one can ask whether a biologically coherent microbial *trait class* is shifted at the sample level.

Bacterial amyloids provide a strong candidate trait class for PD gut microbiome research. Curli fibres are extracellular amyloid structures involved in biofilm formation, adhesion, environmental persistence, and host interaction in *Escherichia coli, Salmonella*, and several related Enterobacteriaceae (Barnhart and Chapman, 2006). Experimental work has connected bacterial amyloid exposure with enhanced alpha-synuclein aggregation in animal and nematode models (Chen et al., 2016), making curli biology relevant to microbiome-centered hypotheses of PD. Importantly, curli biology is not represented by a single bacterial species. A sample may contain multiple low-abundance or moderate-abundance curli-carrier taxa whose combined burden is more informative than any one taxon alone.

This study introduces **Curli Carrier Burden** (CCB), a weighted abundance index that converts a curated curli-carrier taxon panel into a sample-level quantitative trait. CCB does not claim to directly measure curli gene expression or operon completeness. Instead, it provides a conservative, transparent, and reproducible proxy for the processed abundance of bacteria with curli-carrier potential. Its novelty lies in bringing together microbiological curation, explicit confidence weighting, sample-wise trait aggregation, and cohort-wise evidence interpretation into one index.

The purpose of the present work is therefore not to produce a diagnostic classifier or a formal pooled meta-analysis. The purpose is to establish CCB as a mathematically defined and biologically motivated microbial trait index, evaluate its behavior across public processed PD gut metagenomic evidence streams, and define how this index can guide future gene-centric and strain-resolved studies of the curli-vagal PD axis.

## Methods

### Study design

This was a computational, cohort-wise analysis of processed PD gut metagenomic datasets. The analytical workflow had four linked layers: (i) curation of curli-carrier candidate taxa, (ii) construction of a weighted sample-wise CCB index, (iii) cohort-specific PD-control analysis, and (iv) descriptive cross-cohort evidence interpretation. The design emphasizes transparent trait quantification rather than black-box prediction.

**Figure 1:**
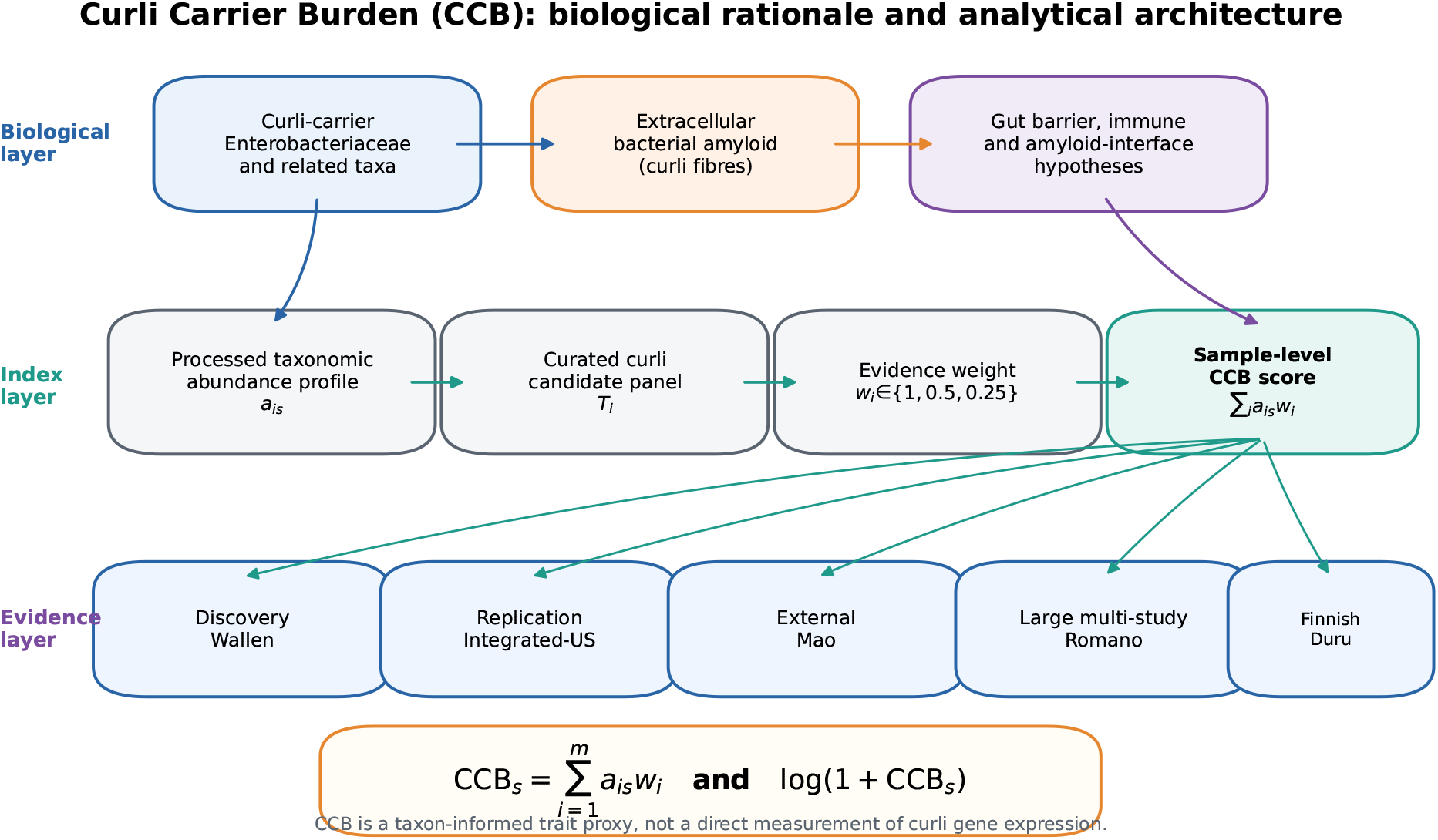
Biological and analytical architecture of Curli Carrier Burden. CCB connects a biological layer of curli-carrier bacterial taxa and bacterial amyloid relevance with a computational layer that combines processed abundance values, curated taxon evidence, and confidence weights. The resulting sample-wise burden score is then evaluated cohort-wise across PD gut metagenomic datasets.

### Processed metagenomic evidence streams

The main analysis focused on five evidence streams: Wallen 2022, Integrated-US, Mao Central China, Romano non-Wallen, and DuruIC 2024. Wallen was treated as the discovery stream. Integrated-US and Mao Central China were treated as replication or external evaluation streams. Romano non-Wallen was analyzed after excluding Wallen-derived samples to reduce discovery-replication overlap. DuruIC 2024 was extracted through the ASAP-MAC/processed metagenomic data route and provided an additional independent Finnish fecal shotgun cohort (Duru et al., 2024; ASAP-MAC, 2026). The cohort summary is shown in Table 1.

**Table 1:** Main processed PD gut metagenomic cohorts used for CCB evaluation.

| Cohort | Analytical role | $n$ | PD | HC/control | Main interpretation role |
| --- | --- | --- | --- | --- | --- |
| Wallen 2022 | Discovery | 724 | 490 | 234 | Initial CCB signal evaluation |
| Integrated-US | Independent support | 244 | 95 | 149 | Supportive processed-cohort replication |
| Mao Central China | External evaluation | 78 | 39 | 39 | Geographically distinct directional evidence |
| Romano non-Wallen | Multi-study external evaluation | 600 | 280 | 320 | Cohort-aware robustness analysis |
| DuruIC 2024 | Additional external evaluation | 134 | 68 | 66 | Independent Finnish processed-cohort test |

**Table 2:** Mathematical components of the Curli Carrier Burden framework.

| Symbol/component | Definition | Interpretation |
| --- | --- | --- |
| $\mathcal{C}$ | Curated candidate set | Evidence-supported taxa with curli-carrier or curli-plausible biology |
| $t_i$ | Candidate taxon $i$ | Species, species-complex, or best available processed taxonomic unit |
| $a_{is}$ | Relative abundance of $t_i$ in sample $s$ | Processed metagenomic abundance value |
| $w_i$ | Evidence weight for $t_i$ | Confidence-adjusted contribution to CCB |
| $\text{CCB}_s$ | $\sum_i a_{is} w_i$ | Sample-wise weighted burden of curli-carrier taxa |
| $\log(1 + \text{CCB})_s$ | $\log(1 + \text{CCB}_s)$ | Scale-stabilized form for distributional comparison |
| $K_d$ | Number of matched candidate taxa in cohort $d$ | Compatibility between processed taxonomy and the CCB panel |

**Table 3:** Cohort-wise quantitative evidence for Curli Carrier Burden across PD gut metagenomic evidence streams.

| Cohort | $n$ | PD | HC | Matched taxa | PD mean | HC mean | PD median | HC median | $p$ value | Evidence class |
| --- | --- | --- | --- | --- | --- | --- | --- | --- | --- | --- |
| Wallen 2022 | 724 | 490 | 234 | 31 | 1.8100 | 1.3900 | 0.0680 | 0.0190 | 0.0020 | Discovery signal |
| Integrated-US | 244 | 95 | 149 | 18 | 2.0000 | 1.8200 | 0.1250 | 0.0000 | 0.0079 | Supportive replication |
| Mao Central China | 78 | 39 | 39 | 17 | 6.8800 | 4.5600 | 2.0800 | 1.7100 | 0.2673 | Directional support |
| Romano non-Wallen | 600 | 280 | 320 | 29 | 0.0153 | 0.0104 | 0.0010 | 0.0006 | 0.0036 | Cohort-sensitive support |
| DuruIC 2024 | 134 | 68 | 66 | 42 | 1.1817 | 1.1663 | 0.1546 | 0.0353 | 0.3069 | Directional support |

### Curli-carrier candidate panel

The CCB candidate panel was constructed from project-curated curli-associated bacterial taxa. The panel included representative curli-carrier or curli-plausible taxa from Enterobacteriaceae and related groups, including *Escherichia, Klebsiella, Enterobacter, Citrobacter, Salmonella, Hafnia, Kluyvera, Leclercia, Kosakonia, Lelliottia*, and related lineages where taxonomic and microbiological evidence supported inclusion. Candidate taxa were matched to processed abundance tables at the most specific available taxonomic resolution. Species-level exact matching was preferred; genus-level or ambiguous species-complex matching was retained only where the processed abundance format limited taxonomic specificity.

### Mathematical definition of Curli Carrier Burden

Let 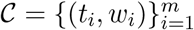 denote the curated curli-carrier candidate set, where *t*_*i*_ is candidate taxon *i* and *w*_*i*_ is its evidence weight. Let *a*_*is*_ be the relative abundance of taxon *t*_*i*_ in sample *s* after matching the curated candidate set to the processed abundance table. The sample-wise Curli Carrier Burden is defined as:

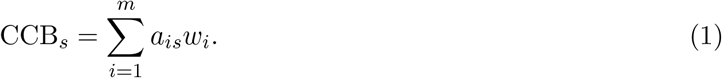

For analyses benefiting from scale stabilization, the transformed burden was defined as:

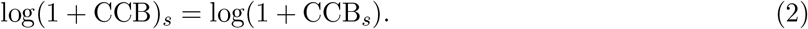

The weighting function was defined as:

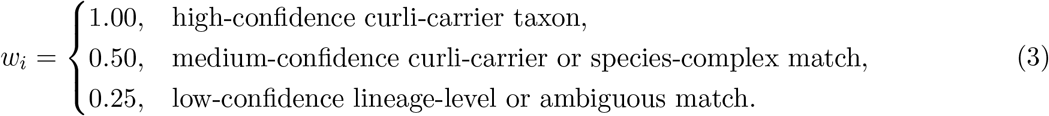

Thus, CCB is a weighted linear burden index. It preserves the direction and magnitude of processed abundance values while preventing lower-confidence matches from contributing as strongly as high-confidence species-level matches. The formula is intentionally simple and auditable: every sample-level score can be decomposed into its taxon-level abundance and evidence-weight components.

### Index compatibility and interpretability

Because processed metagenomic datasets differ in taxonomic representation, CCB interpretation requires recording the number of candidate taxa successfully matched in each cohort. For cohort *d*, we define the matched candidate count:

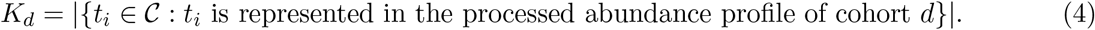

A higher *K*_*d*_ indicates broader compatibility between the processed taxonomic profile and the curated CCB candidate panel. This does not guarantee a positive association with PD, but it improves the interpretability of a cohort as a test of the CCB construct. Cohorts with low matched-taxon coverage remain informative for understanding the boundary conditions of the index, whereas cohorts with moderate-to-high matched-taxon coverage provide stronger tests of aggregate curli-carrier burden.

### Cohort-specific statistical analysis

Within each cohort, PD and control CCB distributions were compared using non-parametric Mann-Whitney U tests. Mean and median CCB were reported to separate distributional shift from outlier-sensitive abundance patterns. Cliff’s *δ* was reported where available as a non-parametric effect-size descriptor. Romano non-Wallen was additionally evaluated using leave-one-study-out analysis, within-study effect profiling, and study-stratified permutation testing. Evidence interpretation used direction, nominal statistical support, effect size, sample size, and matched-taxon compatibility.

### Additional processed-cohort scope checks

Additional ASAP-MAC processed cohorts, including NishiwakiH 2024 and SampsonTR 2025, were explored as scope checks for the CCB framework. These analyses were not used to redefine the index or tune its parameters. Their role was to test how the current CCB panel behaves when processed taxonomic representation differs across cohorts. This is important because CCB is a trait-level index whose interpretability depends on the overlap between the candidate panel and the available taxonomic profiles.

## Results

### CCB converts curli-associated microbiology into a quantitative sample-level trait

The CCB framework transformed a curated list of curli-carrier or curli-plausible bacteria into a sample-level burden score. This is the central contribution of the study. Instead of treating bacterial amyloid biology as a narrative interpretation after taxonomic testing, CCB makes it a measurable variable. Each score has a transparent decomposition into matched taxa, relative abundance values, and evidence weights. This mathematical clarity makes the index reusable, updatable, and comparable across studies.

### Discovery and support across main processed cohorts

Wallen 2022 provided the discovery-stage signal. The analysis included 724 samples, comprising 490 PD and 234 controls, with 31 matched curli-carrier taxa. PD samples showed higher CCB than controls by both mean and median values. PD mean and median CCB were 1.81 and 0.068, respectively, compared with control values of 1.39 and 0.019. The primary Mann-Whitney comparison supported a PD-enriched discovery signal (*p* = 0.0020).

Integrated-US provided supportive independent evidence. This cohort included 244 samples, comprising 95 PD and 149 controls, with 18 matched curli-carrier taxa. PD mean and median CCB were 2.00 and 0.125, respectively, compared with control values of 1.82 and 0.000. The primary Mann-Whitney burden comparison supported higher CCB in PD (*p* = 0.0079).

Mao Central China provided geographically distinct external evidence. The analysis included 78 samples, balanced between 39 PD and 39 controls, with 17 matched burden taxa. PD mean and median CCB were 6.88 and 2.08, respectively, compared with control values of 4.56 and 1.71. The PD-greater-than-control direction was present by both mean and median CCB, although the individual cohort-level comparison was not nominally significant (*p* = 0.2673).

### Romano non-Wallen adds a large multi-study cohort-aware evaluation

The Romano non-Wallen analysis provided a large multi-study external evaluation after excluding Wallen-derived samples. The final set contained 600 samples from 6 studies, including 280 PD and 320 HC samples. The processed mOTUs profile yielded 29 matched curli-associated taxa. PD median CCB was 9.72 × 10^*−*4^, compared with HC median CCB of 5.97 × 10^*−*4^. PD mean CCB was 0.0153, compared with HC mean CCB of 0.0104. The pooled Mann-Whitney test supported higher CCB in PD (*p* = 0.0036), with Cliff’s *δ* = 0.137.

The Romano study-stratified analysis refined this interpretation. Leave-one-study-out analysis retained the PD-greater-than-HC direction by both mean and median CCB. Within-study analyses showed PD-greater-than-HC direction by mean in 4 of 6 studies and by median in 3 of 6 studies. The study-stratified permutation result (*p* = 0.1974) indicates that the Romano signal is best interpreted as cohort-sensitive external support. This is valuable because it demonstrates how CCB can be used not only to detect a pooled signal but also to reveal the cohort structure of that signal.

### DuruIC 2024 provides additional independent directional evidence

DuruIC 2024 added an independent Finnish processed fecal metagenomic evidence stream. The final matched processed set contained 134 samples, comprising 68 PD and 66 HC samples, with 42 matched curli-carrier taxa. PD mean CCB was 1.1817 compared with HC mean CCB of 1.1663. PD median CCB was 0.1546 compared with HC median CCB of 0.0353. The direction was therefore PD-greater-than-HC by both mean and median CCB, although the Mann-Whitney comparison was not nominally significant (*p* = 0.3069, Cliff’s *δ* = 0.1025). DuruIC 2024 therefore supports the value of CCB as a directional, exploratory trait-level index in an independent processed cohort.

**Figure 2:**
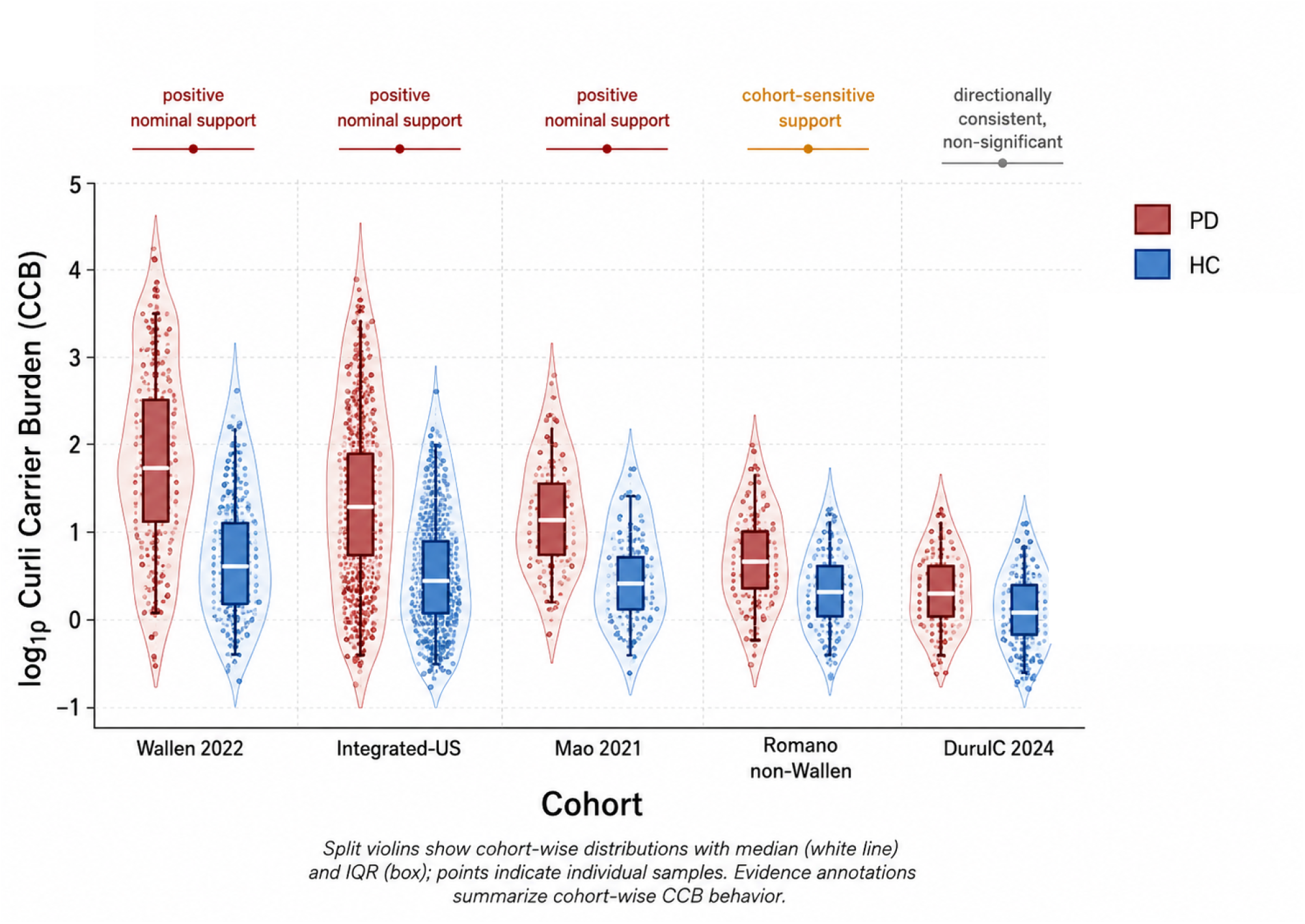
Cohort-wise distributional comparison of Curli Carrier Burden (CCB) across the principal Parkinson’s disease gut microbiome cohorts. Paired violins show PD and HC group distributions within each cohort, with median lines, interquartile boxes, and jittered sample-level points. Evidence annotations above each cohort summarize the cohort-wise CCB behavior without implying formal pooled inference. The figure emphasizes that CCB is an interpretable trait-level burden index whose behavior is recurrent but cohort-dependent across processed metagenomic evidence streams.

**Figure 3:**
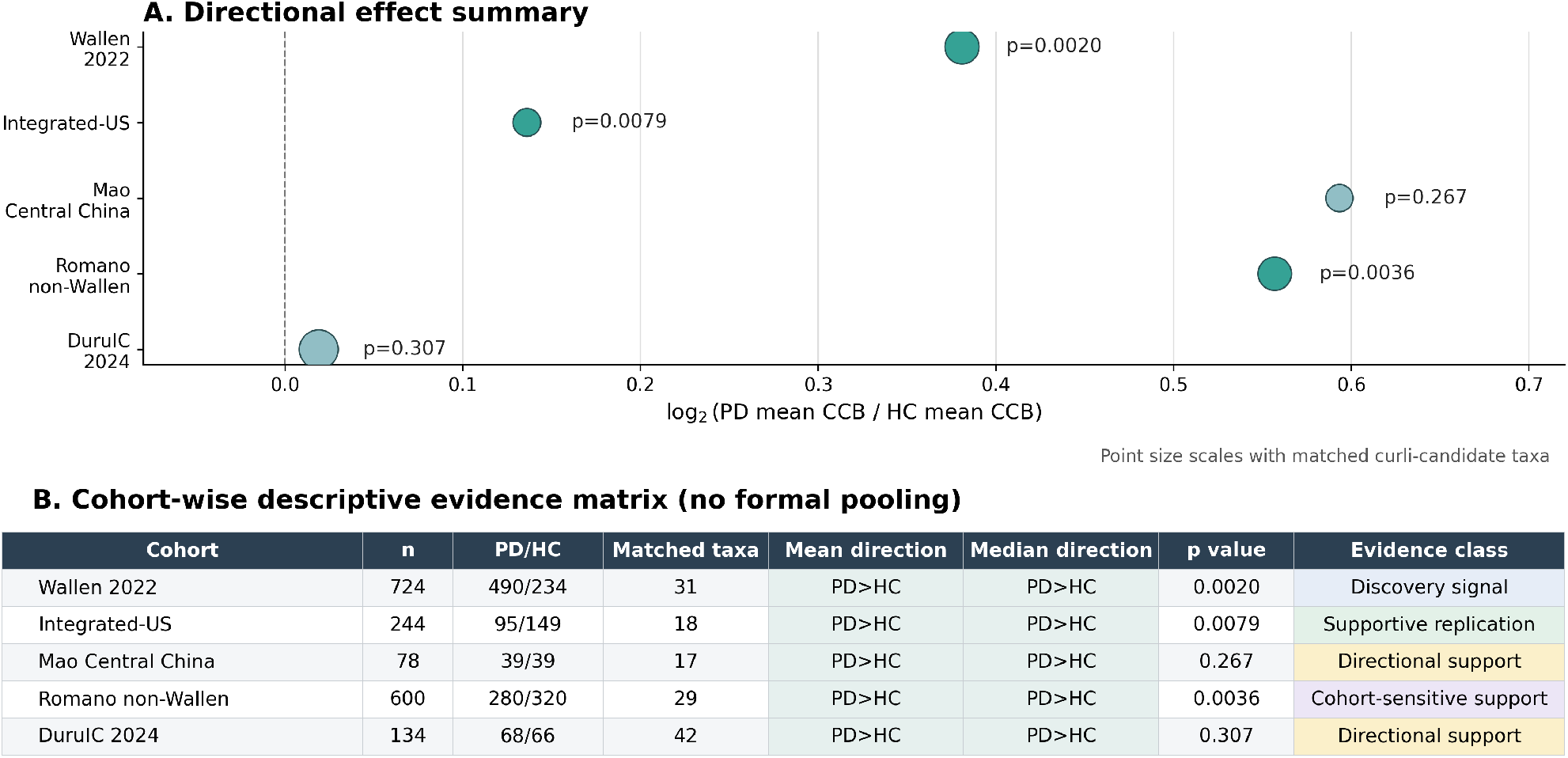
Cohort-wise quantitative evidence summary for CCB. Panel A shows the directional effect as the log_2_ ratio of PD mean CCB to HC mean CCB; point size reflects the number of matched curli-candidate taxa and labels indicate cohort-level Mann-Whitney *p* values. Panel B presents a descriptive evidence matrix summarizing sample size, PD/HC composition, matched-taxon coverage, mean and median direction, nominal *p* value, and evidence class. This figure provides a structured cohort-wise synthesis and is not a formal meta-analysis.

### Processed-cohort scope checks define the boundary conditions of CCB

Additional processed-cohort checks demonstrated the practical importance of taxonomic compatibility. NishiwakiH 2024 produced mixed CCB behavior under the locked processed-data workflow. SampsonTR 2025 was selected a priori as the largest not-yet-analyzed eligible ASAP-MAC cohort, but only one curated curli-carrier taxon was matched in the species-like abundance profile; consequently, it was not a strong test of the full CCB candidate panel. These findings are useful because they show that CCB should always be reported with *K*_*d*_, the number of matched candidate taxa. In this way, the index encourages transparent interpretation instead of forcing all processed cohorts into the same evidentiary category.

## Discussion

This study introduces Curli Carrier Burden as a quantitative trait-level microbiome index for PD gut metagenomic research. The main novelty is not the observation that PD microbiomes differ from controls; that has already been reported by several studies. The novelty is the construction of a microbiology-informed burden variable that aggregates amyloidogenic curli-carrier taxa into a transparent, sample-wise score. This index creates a bridge between taxonomic metagenomic profiles and a biologically meaningful bacterial trait.

The mathematical design of CCB is intentionally simple. It is a weighted abundance sum, but that simplicity is a strength. Every component of the score is interpretable: the candidate taxon, its abundance, and its evidence weight. The framework can therefore be inspected, updated, challenged, and reused. If future experimental or genomic studies refine the curli-carrier status of particular taxa, the candidate panel and weights can be updated without changing the structure of the index. If future datasets provide gene-level csg abundance or strain-level operon completeness, these can be integrated as higher-resolution weights or validation layers.

The cohort-wise results support the utility of CCB as an exploratory microbiome trait index. Wallen and Integrated-US provided nominally supported PD-associated elevation. Mao Central China and DuruIC 2024 showed the same PD-greater-than-HC direction by both mean and median CCB, although their individual cohort-level tests were not nominally significant. Romano non-Wallen added a large multi-study analysis in which pooled CCB was higher in PD and leave-one-study-out direction was retained, while stratified permutation emphasized cohort-sensitive structure. This pattern is biologically meaningful because public PD microbiome cohorts are not interchangeable. They differ in geography, recruitment, sequencing, preprocessing, and taxonomic representation. CCB helps summarize a biologically motivated trait while preserving the ability to inspect cohort dependence.

The additional scope checks further clarify how CCB should be used. A processed cohort with very low matched-taxon coverage is not an equally strong test of the CCB construct. This is not a failure of the biological idea; rather, it is a reminder that trait-level indices depend on how well the trait is represented in the available feature table. The matched-taxon count *K*_*d*_ is therefore an essential companion metric. Reporting *K*_*d*_ alongside CCB prevents overinterpretation and allows future studies to distinguish biological absence from technical or taxonomic under-representation.

The importance of CCB lies in its ability to convert a mechanistic microbiome hypothesis into a measurable feature. Curli biology is relevant to bacterial adhesion, biofilm ecology, host interaction, and bacterial amyloid exposure. These features are difficult to capture using single-taxon differential abundance testing alone. CCB allows investigators to ask a more biologically organized question: does the aggregate burden of curli-carrier taxa differ between PD and controls? This is a different question from whether *E. coli, Klebsiella*, or *Enterobacter* alone is enriched. The index therefore opens a path toward more structured analyses of microbial amyloid ecology.

The present work also provides a foundation for the next generation of curli-vagal PD studies. CCB can be used to prioritize samples for raw-read csgA-csgG mapping, identify cohorts with sufficient curli-candidate coverage, and select high-CCB and low-CCB samples for gene-centric validation. A direct next step is a Curli Operon Integrity Score (COIS), in which read-supported csg operon completeness, csgA/csgG abundance, and strain-level taxonomic assignment are integrated. Such a pipeline would move from taxon-informed exposure potential to gene-supported curli capacity. This is precisely where the current index is most useful: it provides a rational discovery layer that can guide deeper, more expensive validation.

The framework can also support subtype-aware PD research. Gastrointestinal symptoms, constipation, rapid eye movement sleep behavior disorder, hyposmia, autonomic features, and body-first/vagal hypotheses are all relevant to PD heterogeneity. Once raw-read csg evidence and richer clinical metadata are available, CCB and COIS can be evaluated against these phenotypic axes. This would allow future studies to ask whether microbial amyloid burden is associated with particular PD subgroups rather than with PD status alone.

**Figure 4:**
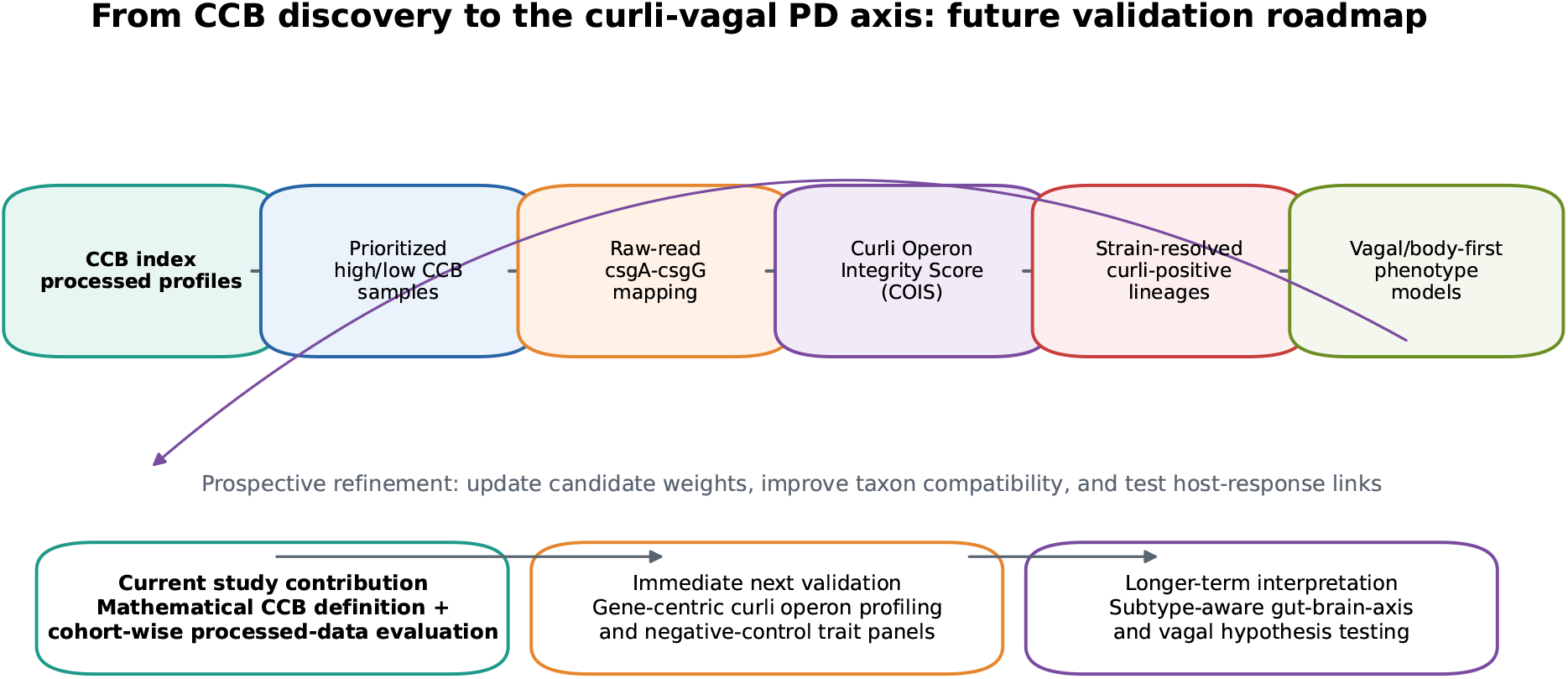
Future validation roadmap from CCB to the curli-vagal PD axis. The current study establishes the mathematical CCB index and evaluates it across processed metagenomic cohorts. The next analytical layer can use CCB to prioritize raw-read csgA-csgG mapping, Curli Operon Integrity Score construction, strain-resolved curli-positive lineage analysis, and subtype-aware gut-brain-axis modeling.

Overall, CCB should be viewed as a reusable analytical framework rather than a final mechanistic endpoint. Its value is that it makes bacterial amyloid carrier burden measurable in public processed metagenomic datasets and creates a structured hypothesis-generating bridge to gene-centric and strain-resolved validation. The current results support the biological plausibility and practical utility of this approach while also showing why future work should explicitly report taxon compatibility, cohort structure, and raw-read validation.

## Conclusions

We introduced Curli Carrier Burden as a novel, weighted, taxon-informed index for estimating amyloidogenic curli-carrier bacterial burden in processed gut metagenomic profiles. Across main PD gut microbiome evidence streams, CCB showed PD-associated elevation in several cohorts and directionally consistent behavior in additional independent data, with cohort-aware analyses highlighting the importance of matched-taxon compatibility and study structure. The framework provides a clear mathematical foundation for future microbiome trait-burden studies and establishes a practical route toward raw-read csg operon validation, strain-resolved curli biology, and deeper investigation of the curli-vagal PD axis.

## Declarations

### Ethics approval and consent to participate

This study used public or processed de-identified metagenomic data resources and did not involve new human participant recruitment, sample collection, or wet-laboratory experimentation.

### Consent for publication

Not applicable.

### Availability of data and materials

The analysis used publicly available or processed metagenomic resources and derived project reports. The analysis code, smoke tests, non-sensitive summary reports, figure-generation resources, and manuscript-supporting files are publicly available at GitHub: https://github.com/drnabanitaghosh888-de/curli-carrier-burden-pd. The version of record used for manuscript preparation has been archived in Zenodo with DOI: 10.5281/zenodo.20371789. Controlled-access datasets, raw sequencing reads, and participant-level restricted metadata are not redistributed and should be obtained from the original repositories according to their respective access conditions and data-use agreements.

### Authors’ contributions

NG conceptualized the study. NG led the biological framing of the curli-vagal PD hypothesis, contributed to interpretation of bacterial amyloid relevance, and critically shaped the manuscript. NG and KS designed and implemented the computational CCB framework, performed the cohort-wise analyses, generated figures and tables, and contributed to manuscript drafting and revision. Both authors read and approved the final version.

## Acknowledgements

The authors acknowledge the investigators and data-resource teams who generated, curated, and released the Parkinson’s disease gut metagenomic datasets and processed resources analyzed in this work.

## References

Ghosh N, Sinha K. Curli Carrier Burden analysis for Parkinson’s disease gut microbiomes. Zenodo.2026. doi:10.5281/zenodo.20371789. Available at: https://doi.org/10.5281/zenodo.20371789.

ASAP-MAC. parkinsonsMetagenomicData: uniformly processed gut microbiome data from Parkinson’s disease studies.GitHub repository. Available at: https://github.com/ASAP-MAC/parkinsonsMetagenomicData. Accessed 2026.

Barnhart MM, Chapman MR. Curli biogenesis and function. Annual Review of Microbiology. 2006;60:131–147. doi:10.1146/annurev.micro.60.080805.142106.

Bedarf JR, Hildebrand F, Coelho LP, Sunagawa S, Bahram M, Goeser F, Bork P, Wullner U. Functional implications of microbial and viral gut metagenome changes in early stage L-DOPA-naive Parkinson’s disease patients. Genome Medicine. 2017;9:39. doi:10.1186/s13073-017-0428-y.

Boktor JC, Sharon G, Verhagen Metman LA, Lee SJ, Matheoud D, Sampson TR, Gradinaru V. Integrated multi-cohort analysis of the Parkinson’s disease gut metagenome. Movement Disorders. 2023;38:399–409. doi:10.1002/mds.29300.

Chen SG, Stribinskis V, Rane MJ, Demuth DR, Gozal E, Roberts AM, Jagadapillai R, Liu R, Choe KY, Shivakumar B, Son F, Jin S, Kerber R, Adame A, Masliah E, Friedland RP. Exposure to the functional bacterial amyloid protein curli enhances alpha-synuclein aggregation in aged Fischer 344 rats and Caenorhabditis elegans. Scientific Reports. 2016;6:34477. doi:10.1038/srep34477.

Duru IC, Lecomte A, Shishido TK, et al. Metagenome-assembled microbial genomes from Parkinson’s disease fecal samples. Scientific Reports. 2024;14:18906. doi:10.1038/s41598-024-69742-4.

Mao L, Zhang Q, Li H, Chen Y, Li Y, Huang X, Zheng P, Li J, Li X, Chen J, Xie P. Cross-sectional study on the gut microbiome of Parkinson’s disease patients in central China. Frontiers in Microbiology. 2021;12:728479. doi:10.3389/fmicb.2021.728479.

Nishiwaki H, Ito M, Ishida T, et al. Meta-analysis of shotgun sequencing of gut microbiota in Parkinson’s disease. npj Parkinson’s Disease. 2024. doi:10.1038/s41531-024-00724-z.

Pasolli E, Schiffer L, Manghi P, et al. Accessible, curated metagenomic data through ExperimentHub.Nature Methods. 2017;14:1023–1024. doi:10.1038/nmeth.4468.

Qian Y, Yang X, Xu S, Huang P, Li B, Du J, He Y, Su B, Xu L, Wu B, Wang L, Liu M, Chen S, Wang X, Jin H, Zhang L, Li H, Zhang Y, Liu J, Wang H. Gut metagenomics-derived genes as potential biomarkers of Parkinson’s disease. Brain. 2020;143:2474–2489. doi:10.1093/brain/awaa201.

Romano S, Savva GM, Bedarf JR, Charles IG, Hildebrand F, Narbad A. Meta-analysis of the Parkinson’s disease gut microbiome suggests alterations linked to intestinal inflammation. npj Parkinson’s Disease. 2021;7:27. doi:10.1038/s41531-021-00156-z.

Wallen ZD, Appah M, Dean MN, Sesler CL, Factor SA, Molho E, Zabetian CP, Standaert DG, Payami H. Metagenomics of Parkinson’s disease implicates the gut microbiome in multiple disease mechanisms.Nature Communications. 2022;13:6958. doi:10.1038/s41467-022-34667-x.

